# A Unified Architecture of Transcriptional Regulatory Elements

**DOI:** 10.1101/019844

**Authors:** Robin Andersson, Albin Sandelin, Charles G. Danko

## Abstract

Gene expression is precisely controlled in time and space through the integration of signals that act at gene promoters and gene-distal enhancers. Classically, promoters and enhancers are considered separate classes of regulatory elements, often distinguished by histone modifications. However, recent studies have revealed broad similarities between enhancers and promoters, blurring the distinction: active enhancers often initiate transcription, and some gene promoters have the potential of enhancing transcriptional output of other promoters. Here, we propose a model in which promoters and enhancers are considered a single class of functional element, with a unified architecture for transcription initiation. The context of interacting regulatory elements, and surrounding sequences, determine local transcriptional output as well as the enhancer and promoter activities of individual elements.

## The classical view: enhancers and promoters as distinct regulatory elements

Transcriptional regulatory DNA sequences encode intricate programs of gene expression that are responsible for controlling virtually all cellular functions. These regulatory sequences recruit proteins known as transcription factors (TFs) in a DNA sequence-dependent fashion, allowing cells to precisely control the rates of chromatin decompaction, transcription initiation, and the release of RNA polymerase II (RNAPII) into productive elongation [1,2]. Classically, a sharp distinction has been drawn between two types of regulatory sequence, promoters and enhancers.

Promoters are DNA sequences that regulate and initiate RNAPII transcription at proximal transcription start sites (TSS). RNAPII core promoters can encompass as little as 100bp of DNA surrounding the TSS, which often contains one or more degenerate copies of the core DNA sequence elements, including the TATA-box and the Initiator (INR). These core promoter elements are recognized by general transcription factors (GTFs) such as TFIID and TFIIB, which are responsible for recruiting and assembling the RNAPII pre-initiation complex (PIC) [3]. RNAPII PIC assembly and transcription initiation are further facilitated by TFs bound proximally to core promoters. While promoters were originally found at TSSs of known genes, more direct experimental methods, such as sequencing 5’ ends of RNAs, have identified promoters genome-wide and showed that a much larger proportion of mammalian genomes is associated with transcription initiation than what can be accounted for by annotated gene models [4,5]. Promoters, here referring to units of proximal and core promoter regions collectively, work together with other regulatory regions, like enhancers and silencers, to regulate all stages of RNAPII transcription from RNAPII recruitment to transcriptional elongation.

Enhancers, in contrast to gene promoters, are generally considered TF-binding regulatory regions distal to gene TSSs. The first enhancer discovered was a 72 bp tandem repeat upstream of early genes in the simian virus 40 (SV40) genome [6,7]. This sequence was reported to increase transcription of the β-globin gene more than 200-fold when inserted into the same recombinant expression vector, irrespective of its position, distance, and orientation relative to the target gene promoter [8]. Importantly, transcription of the β-globin gene invariably initiated at the β-globin promoter, indicating that the enhancer worked to stimulate transcription from the target promoter at a distance. Enhancers were shortly thereafter discovered in the mouse genome [9] and are today considered key players of transcriptional regulation across Eukaryota. Although different models have been proposed to explain how enhancers regulate gene expression over long genomic distances (Box I), several recent studies strongly suggest that chromatin architectures place enhancers in close three-dimensional proximity with target gene promoters [10]. Based on these observations, enhancers are classically defined by their ability to increase transcriptional output from target genes.

However, recent studies are challenging the conventional wisdom that enhancers and promoters are distinct entities. These results are changing the way that regulatory elements are defined and identified. Here, we review recent progress in the area, focusing on new genome-wide studies reporting broad similarities in the biochemical and DNA sequence properties of mammalian enhancers and gene promoters. Based on these new findings, we argue here that the discriminatory view of enhancers and promoters as distinct regulatory elements is problematic because it suggests that they have distinct functions. Rather, their broad similarities and overlapping functional properties suggest a unifying view of enhancers and promoters as a single class of ‘regulatory element’, each with an intrinsic ability to drive local transcription (i.e., act as a ‘promoter’) or enhance distal transcription (i.e., act as an ‘enhancer’) with varying strengths, and that their primary function is likely context-dependent.

#### Box I: Models for enhancer function

Several models have been proposed for how enhancers regulate expression at gene promoters over long genomic distances, in some cases up to a megabase in units of linear DNA. A popular hypothesis is that enhancers regulate transcription by looping into close three-dimensional proximity with target gene promoters [66]. This model is supported by the frequencies of distal DNA sequence ligation using chromatin conformation capture techniques as well as by florescence in situ hybridization (FISH) [67]. However, while 3C results indicate physical interactions between enhancers and promoters, specific interactions cannot always be reproduced using FISH [68], suggesting that technical work still needs to be done before we are able to reproducibly capture interactions between enhancers and promoters. Other models have also been proposed that explain how enhancers regulate gene expression, including promoter tracking [69] and enhancer-promoter linking via protein bridges [70], and these are generally not mutually exclusive with DNA looping. In addition to the constraints on transcriptional regulation posed by larger chromatin architectures, context-dependent properties influence whether physical proximity of regulatory elements lead to increased transcriptional output, including enhancer specificity for certain core promoter elements [44,71], promoter competition [72], and insulation [73].

Little is known about the molecular mechanisms by which enhancers regulate the transcription level at target gene promoters. Several mechanisms have been proposed, including i) facilitating or enabling transcription initiation by supplying needed factors like the Mediator [74], GTFs [27] or RNAPII [23], ii) affecting the release rate of RNAPII into productive elongation by activation of P-TEFb [75], or iii) by recruitment of the super elongation complex [76] (for a recent review of enhancer function see [2]).

## The classical approach: classification of regulatory elements by histone modifications

Several studies have revealed post-translational modifications of chromatin that are characteristic of regulatory sequences across the genome (Box II) [11]. Certain marks have genomic distributions that correlate broadly with the expected location of either enhancers or gene promoters. Most notably, a seminal study found that H3K4me3 is highly enriched in regions that are proximal to the 5’ end of gene annotations where active gene promoters are expected to reside, whereas H3K4me1 is more frequently found distal to gene promoters [12]. This led to the suggestion that H3K4me3 and H3K4me1 could discriminate between promoters and enhancers. As these modifications are not mutually exclusive, their signal ratio (H3K4me1:H3K4me3) – found to be low at promoters and high at enhancers – was suggested to aid in discrimination [13]. In addition, H3K27ac has been proposed to distinguish active enhancers and promoters from inactive ones [14,15]. In the past few years, classification of regulatory sequences based on histone modifications have become widely adopted by the genomics and gene regulation communities.

## Active enhancers can independently work as promoters

Recent observations point to numerous functional similarities between enhancers and gene promoters. RNAPII binds and initiates transcription at active enhancers [16,17], demonstrating that at least some enhancers direct the biological process that is a defining feature of gene promoters. More recently, it has become clear that tens of thousands of enhancers initiate RNAPII in opposing orientations from local TSSs and transcribe enhancer-templated non-coding RNAs (eRNAs), albeit in many cases with a lower rate of transcription initiation than at gene promoters [18-22] (Figure 1A). Although RNAPII could be partially supplied to enhancers by gene promoter regions that are physically proximal, at least some enhancers are able to initiate transcription independently of gene promoters. For example, GTF and RNAPII recruitment to enhancers upstream of the α-globin gene precedes promoter recruitment, and is unaffected in MEL cell line hybrids lacking the α-globin gene promoter [23]. Another example of the dual role of enhancers with local promoter activities are a group of α-globin enhancers contained within *Nprl3* gene, which act as both enhancers to the α-globin gene promoter as well as alternative promoters to non-coding RNAs sharing exons with *Nprl3* [24]. Like active gene promoters, active enhancers are demarcated by a well-positioned array of nucleosomes surrounding a nucleosome-depleted region (NDR) [21,25,26], and the distance between divergent TSSs and NDR edges is constrained [19,21,25]. Consistent with their ability to recruit RNAPII, enhancer NDRs are enriched in typical core promoter sites, including TATA and INR motifs, and bind GTFs [19,22,27,28] (Figure 1A). In general, enhancers are depleted of CpG islands and seem to recruit similar repertoires of lineage-specific master regulator TFs as CpG-poor gene promoters [21]. Taken together, these observations strongly suggest that enhancers can independently work as promoters, and are similar to gene promoters in terms of DNA sequence, nucleosome positioning, and TF binding.

**Figure 1:**
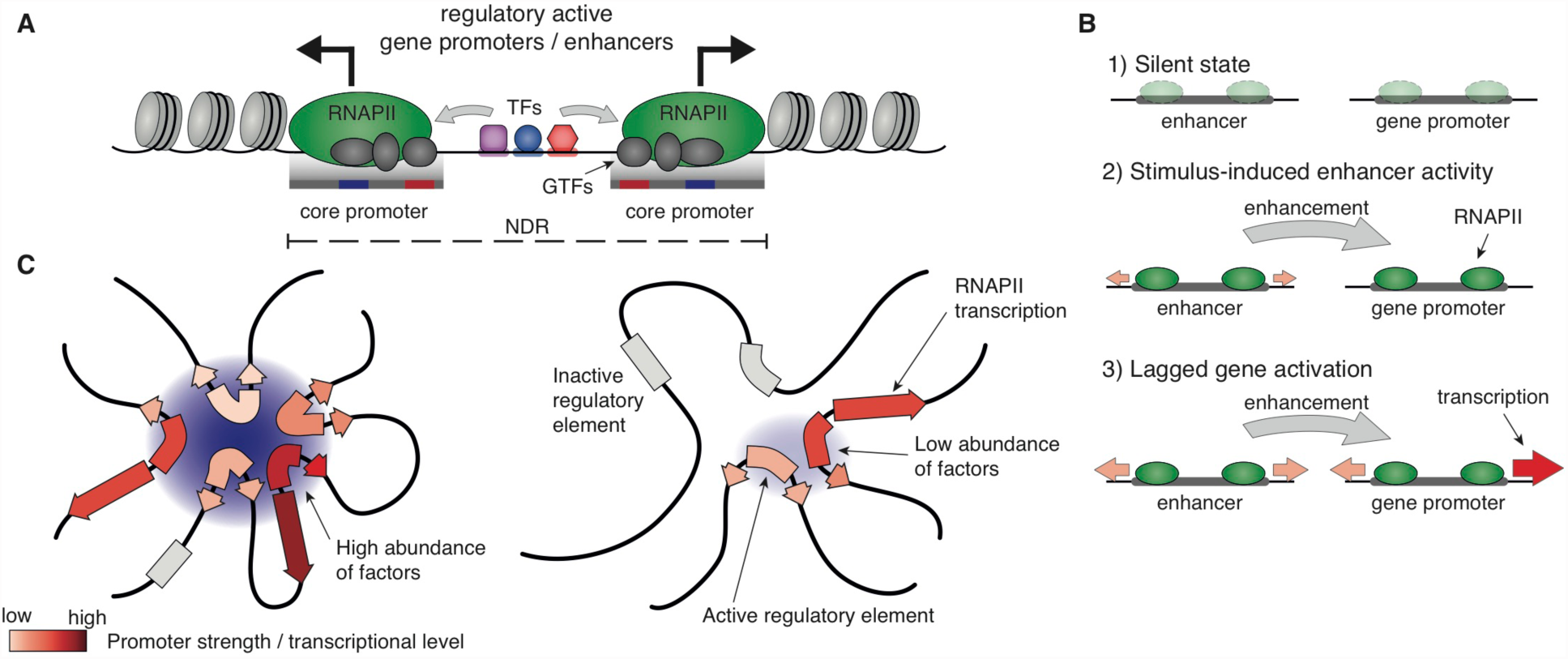
Active regulatory elements are divergently transcribed. **A**: Both regulatory active gene promoters and gene-distal enhancers are transcribed. RNA polymerase II (RNAPII) recruitment and transcription initiation are mediated by general transcription factors (GTFs) binding core promoter regions in close proximity to flanking nucleosomes. This is facilitated by transcription factors (TFs), which often bind proximal to core promoters. Transcription often initiates divergently and at the boundary of the nucleosome-depleted region (NDR). **B**: Gene expression is often preceded by, or changes concurrently with, changes in enhancer transcription. In a silent (non-expressed) state (1), enhancers and promoters may, or may not, bind RNAPII. Upon stimulus (2), transcriptional activity at enhancers marks regulatory enhancer activity with local transcription and increases in RNAPII recruitment at the target gene promoter. (3) Gene expression may lag behind transcriptional activation at enhancers. **C**: Chromatin interactions place regulatory elements in close physical proximity. The individual properties of regulatory elements (chromatin characteristics as well as TF and RNAPII recruitment strengths) as well as context-dependent properties (such as promoter competition, insulation, and core promoter specificity) jointly determine the formation of multiple regulatory interactions (Box I). Via regulatory cooperation, multiple regulatory elements may increase the local concentration of factors (TFs, GTFs, co-activators, and RNAPII) needed for transcription in RNAPII-enriched foci (left panel) and thereby achieve in aggregate different levels of transcriptional activity than of RNAPII foci including fewer regulatory elements (right panel). Nucleosome illustrations in panel A are reused with permission from [38]. Panel C is modified with permission from [38].

#### Box II: Regulatory elements and associated chromatin modification states

The N-terminal tails of histone proteins H2A, H2B, H3, and H4 within nucleosomes can be chemically modified. In particular, amino acid residues can be methylated or acetylated, which correlates with functional properties of the chromatin. For instance, H3K27ac (acetylation of lysine 27 in histone H3) is correlated with transcriptional activity. The locations of histones “marked” with a certain histone modification can be measured genome-wide with chromatin immunoprecipitation (ChIP) coupled with either tiling microarrays (ChIP-chip) or sequencing (ChIP-seq) [38]. Profiling of histone modifications in different cell types have enabled systematic inference of different types of functional genomic entities [12,53,77,78]. Such studies have shown that, typically, the nucleosomes flanking active gene promoters are enriched in H3K27ac, H3K4me3 (trimethylation of lysine 4 in histone H3), and to some degree H3K4me1 (mono-methylation), whereas enhancers are enriched in H3K27ac and H3K4me1 but not H3K4me3, or relatively low levels thereof. In addition, several marks have been found to localize at both enhancers and promoters. Most notably, H3K27ac marks active promoters and a subset of enhancers that appear to be actively regulating gene transcription, leading to the notion of classifying regulatory elements into ‘active’ and ‘poised’ classes [15,79]. Poised regulatory elements sometimes carry bivalent marks associated with activity and repression at the same time, for instance promoters having H3K4me3 and H3K27me3 (the latter mark is most often be found at inactive gene promoters). Collectively, these studies have argued that functional elements can be effectively annotated by their distinct patterns of histone modifications.

Although many enhancers are transcribed, they represent a subset of enhancers predicted from histone modifications. However, untranscribed enhancers in one cell type are often transcribed in another [21,25], suggesting that many enhancers have promoter activities in an appropriate context. More generally, this observation raises the question of what makes transcribed enhancers different from untranscribed ones? Although this question is unresolved, a few key observations have been made. Mammalian enhancers producing eRNAs are more likely to interact with gene promoters [29] and validate by *in vitro* reporter gene assays [21,30], compared to untranscribed enhancers identified using histone marks. Furthermore, using a transgenic mouse enhancer assay, tissue specificity of eRNA expression was shown to correctly predict tissue specific *in vivo* enhancer activity of 88% of known enhancers that were tested [31]. These results suggest that the promoter activity of an enhancer across cells is a proxy of its enhancer activity. This conclusion is also supported by the observation that transcription at enhancers often changes in a stimulus-dependent manner [32-36] and correlates with changes in the transcriptional output of target genes [18,21], often with dynamics that match or precede gene activation [20,33,35] (Figure 1B).

## Gene promoters possess enhancer potential

Since active enhancers have promoter activities, it follows that what we classically refer to as promoters could also function as enhancers in certain situations. Although in general a neglected topic of research, we find some clues in studies of larger chromatin architectures.

Regulatory elements often interact in close physical proximity in RNAPII foci that are sometimes referred to as transcription factories [37] and the activity of each regulatory element within these foci can be influenced by other regulatory elements involved (Figure 1C). Although the regulatory mechanisms remain poorly characterized, a common model posits that regulatory sequences synergistically achieve high levels of transcriptional activity by increasing the local concentration of factors needed for transcription [37-41] (Figure 1C). Studies based on chromatin conformation capture have observed RNAPII foci involving multiple gene promoters, with few or no enhancers [29,42]. In such arrangements, the degree to which each gene promoter acts as a promoter or enhancer may be dependent on the local context of chromatin [42], expressed TFs, and sequence context (e.g., the occurrence and strength of core promoter elements and TF binding sites). Li *et al.* found certain gene promoters in such constellations in MCF7 cells to possess enhancer potential using *in vitro* reporter assays [42], suggesting that they might enhance the transcriptional activity of promoters to which they are physically proximal. In line with these results, Leung *et al.* predicted a considerable fraction of strong promoters in one tissue as enhancers in other tissues and demonstrated both activities *in vitro* [43]. Finally, by measuring the enhancer potential of randomly fragmented genomic DNA using STARR-seq, Zabidi *et al.*, found that a significant number of Drosophila gene promoters have enhancer potential [44]. Taken together, these results suggest that many gene promoters could also act as enhancers. However, a common weakness of these experiments is that they are based on an artificial reporter gene system. Systematic experiments removing DNA sequences that encode promoters will be required to conclusively demonstrate a dual role *in vivo* (Box III). Yet, in spite of this weakness, these findings point toward a dual role of many promoters as transcriptional enhancers.

## Differential RNA stability defines two classes of regulatory sequence

Although gene promoters and gene-distal enhancers have many similarities, the properties of RNAs that they produce differ substantially [21,40,45]. mRNAs are generally multi-exonic, highly abundant, polyadenylated, and exported to the polyribosome allowing translation. By contrast, eRNAs are often non-spliced, non-polyadenylated transcripts that are observed in low copy numbers and retained in the nucleus. In fact, these properties make eRNAs highly similar, if not identical, to antisense transcripts arising upstream from protein-coding gene promoters (uaRNAs/PROMPTs) [40,46,47] (Figure 2). There is now strong evidence that differences between RNA classes are, to a large degree, encoded in DNA sequences around the 5’ end of each transcript. Differences are established after RNAPII initiation and controlled by the location, order, and number of DNA sequence elements [19,46-49]. Utilization of early polyadenylation sites, a discerning characteristic of eRNA and uaRNA/PROMPT transcription, leads to nuclear degradation of RNAs by the exosome complex [21,40,47,50,51] (Figure 2). In contrast, protein-coding transcripts are enriched for TSS-proximal U1 snRNP recognition sites (5’ splice sites) and depleted for early polyadenylation sites. Early U1 binding seems to counteract termination at polyadenylation sites [47-49]. Differences in RNA stability and length reflect the class of the resulting transcript, such that mRNAs most often originate from NDRs encoding stable RNAs, whereas enhancer transcription results in unstable, short RNAs. It follows that a natural and biologically meaningful way to discriminate between gene promoters and gene-distal enhancers, if such discrimination is desired, is by the biochemical properties (including length, splicing events, and stability) and abundances of the RNAs that result from their activation, rather than their perceived regulatory activity.

**Figure 2:**
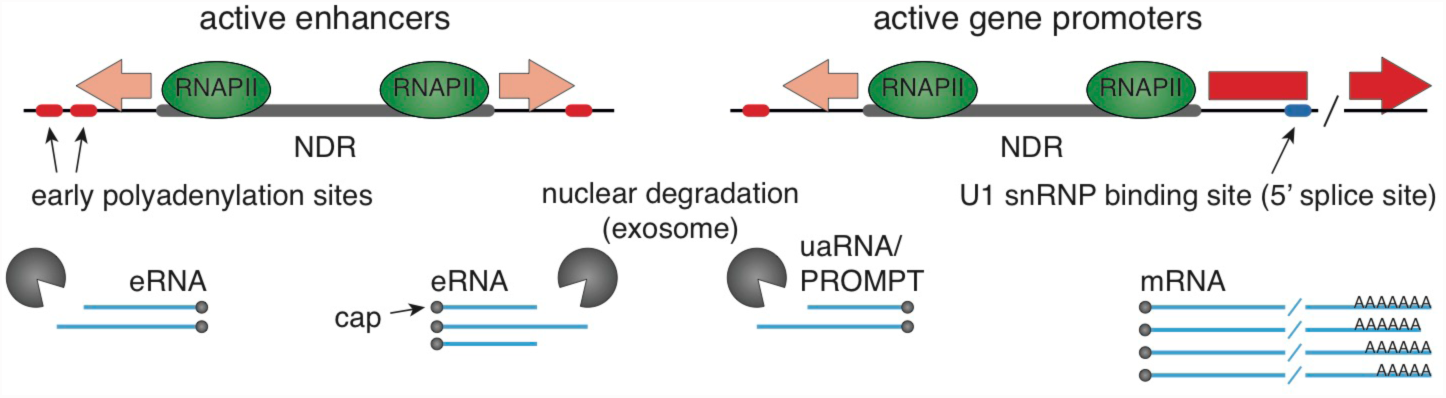
Differential RNA stability distinguishes between regulatory elements. Transcription initiation at gene promoters and gene-distal enhancers results in characteristically different RNAs. eRNAs are typically short, non-spliced and degraded by the nuclear exosome complex, making them highly similar to antisense transcripts (uaRNAs/PROMPTs) produced upstream of mRNA TSSs. The length and instability of eRNAs and uaRNAs/PROMPTs are likely a consequence of RNAPII utilization of early polyadenylation sites. Compared to sequences downstream of eRNA and uaRNA/PROMPT 5’ ends, mRNA transcripts have a decreased occurrence and utilization of early polyadenylation sites while being enriched for TSS-proximal U1 snRNP recognition sites (5’ splice sites). Early U1 binding seems to counteract termination at early polyadenylation sites, which otherwise leads to nuclear degradation of RNAs by the exosome. Consequently, mRNAs are often comparably longer, spliced, and more stable than eRNAs and uaRNAs/PROMPTs and are exported to the polyribosome.

## Differences in chromatin status reflect transcriptional activity, not element type

Related to the apparent overlap in functionality of enhancer and promoters, recent studies have identified numerous exceptions to the general rules proposed for distinguishing enhancers from promoters based on histone modifications. Genome-wide profiling of histone modifications in CD4+ T cells identified an enrichment of H3K4me3 at enhancers [14,52], despite earlier reports that this mark is usually promoter specific. In fact, enhancers marked by H3K4me3 seem to posses high enhancer activity [22,53,54]. Likewise, gene promoters driving lower levels of transcription carry H3K4me3 less frequently, and can often be marked by H3K4me1 [52]. Changes in the ratio between H3K4me1 and H3K4me3 are also commonly observed. Thymocyte enhancers pre-marked with H3K4me1 acquire H3K4me3 upon T-cell receptor induced differentiation [54]. Likewise, relative changes between H3K4me3 and H3K4me1 were discovered following differentiation of H1 embryonic stem cells into a number of early embryonic lineages [43]. Thus, the ratio between H3K4me1 and H3K4me3 at regulatory elements, although usually low at gene promoters and high at enhancers, is a dynamic property, which varies substantially between tissues and biological conditions (Figure 3A). Like H3K4me3, H3K27ac is neither restricted to enhancer nor promoters. Although generally used to distinguish active from inactive enhancers [15], H3K27ac is often observed also at active gene promoters [14,53], to which it has a strong preference [55].

**Figure 3:**
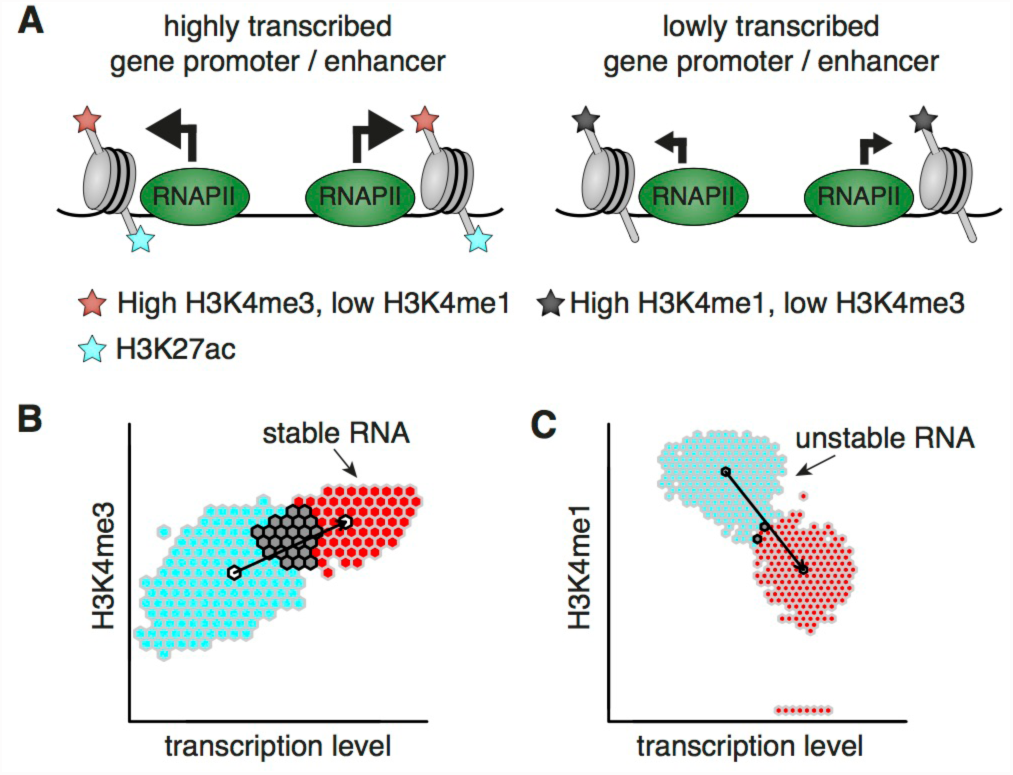
Transcriptional level is related to histone modifications at regulatory elements. **A**: Highly transcribed enhancers are often marked by H3K4me3 and H3K27ac, making them hard to distinguish from transcribed gene promoters using histone modifications alone. Lowly transcribed gene promoters and gene-distal enhancers are rarely marked by, or only by low levels of, H3K4me3 or H3K27ac, but often by H3K4me1. **B**: The level of H3K4me3 at the nucleosome-depleted regions (NDRs) at the 5’ end of transcripts encoding stable RNAs (e.g., mRNA) and unstable RNAs (e.g., eRNA) is correlated with their transcriptional activity (see Figure 2 for the connection between RNA stability and regulatory element) **C**: The level of H3K4me1 at the NDRs of stable RNAs and unstable RNAs is inversely correlated with their transcriptional activity. Nucleosome illustrations in panel A are reused with permission from [38]. Panels B and C are modified with permission from [19].

What, then, is the biological property reflected in these marks? Recent work proposes that histone modifications actually reflect the local level of transcriptional activation driven by each regulatory element. Core *et al.* [19] found that the level of RNAPII, as defined by precision run on sequencing of nascent RNA (PRO-seq), generally increases proportionally with H3K4me3 (Figure 3B) and decreases with H3K4me1 (Figure 3C) at both putative enhancers and promoters. PRO-seq near mammalian promoters reflects the joint contribution of initiation rates and the release rate of RNAPII from pause [56]. Thus, although there is substantial variation between histone methylation and RNAPII levels, it is possible that direct measurements of either initiation or pause release may provide a more direct correlation. This implies that highly transcribed enhancers are marked with high levels of H3K4me3, whereas lowly transcribed or silent gene promoters are marked predominantly by H3K4me1, both predictions which have now been observed in many instances [19,43,54]. Thus, although typical examples of promoters and enhancers carry the H3K4 methylation pattern that are conventionally attributed to them (low versus high H3K4me1:H3K4me3 ratio, respectively), this may simply reflect the lower levels of RNAPII recruited by most enhancers compared with gene promoters. This hypothesis is supported by high relative importance of H3K4me3 as well as H3K27ac in predicting gene expression levels from histone modifications [57].

While the idea that H3K4me3 is linked to transcriptional levels is difficult to test directly, there is some supporting evidence already in the literature. For example, Pekowska *et al.* replaced the endogenous *Tcrb* enhancer with a mutated copy that confers a lower activity and observed a local increase in H3K4me1 and decrease in H3K4me3 compared to wild type [54], apparently supporting a causal relationship between transcriptional activity and histone methylation. This notion is also consistent with reports that H3K4 methyltransferases are recruited by the carboxy-terminal domain of RNAPII [58-60]. Thus, we argue that there is a relationship between H3K4 methylation and transcription levels that applies at both enhancers and gene promoters.

It should be noted that the level of H3K4me3 is not related solely to transcription level, which is reflected by their relatively weak correlation (Figure 3B). While H3K4 Set1 methyltransferases are directly acquired by RNAPII, there is an observed bias of H3K4me3 to CpG-rich sequences mediated by the CpG binding Cfp1 subunit of Set1 complexes [61]. The localization and level of H3K4me3 is further affected by the presence and position of the first exon 5’ splice site, and inhibition of splicing reduces H3K4me3 levels, suggesting a link between splicing and H3K4me3 [62]. However, interfering with splicing is also likely to decrease the nuclear stability of a transcript [46,47], as discussed above, and thus is likely to decrease RNAPII at each regulatory element.

## Understanding the relationship between enhancer and promoter function

The relationship between the potential of each regulatory element to act as a local promoter and its ability to enhance transcription at distal promoters remains poorly understood. Although based upon only a few cases, Li *et al.* observed the strongest *in vitro* luciferase reporter activity for gene promoters with low transcriptional activities and enhancer-like histone modifications [42]. These observations led to a proposed model in which the local enhancer and promoter potentials of a regulatory element follow an inverse relationship: the stronger the enhancer potential of a regulatory element, the weaker its promoter activity [42] (Figure 4A). However, this hypothesis remains to be directly tested. Moreover, it appears to contradict the correlation between enhancer transcription and target gene transcription [20,33,35], which potentially implies that increases in the ability of a regulatory element to enhance RNAPII transcription distally also increases its local promoter-like transcriptional activity (Figure 4B). Yet, although the transcriptional activity of an enhancer seems to be highly predictive of its ability to enhance distal gene expression (enhancer activity), *in vitro* reporter validations did not show any clear relationship between the level of *in vivo* expression of transcribed enhancers and the amount of enhancer activity (measured as the increase in reporter signal enabled by the enhancer) [21]. This might suggest a binary rather than a quantitative relationship between enhancer activity and enhancer expression. Taken together, the relationship between ‘enhancer’ and ‘promoter’ function of each regulatory sequence is likely to be complicated.

**Figure 4:**
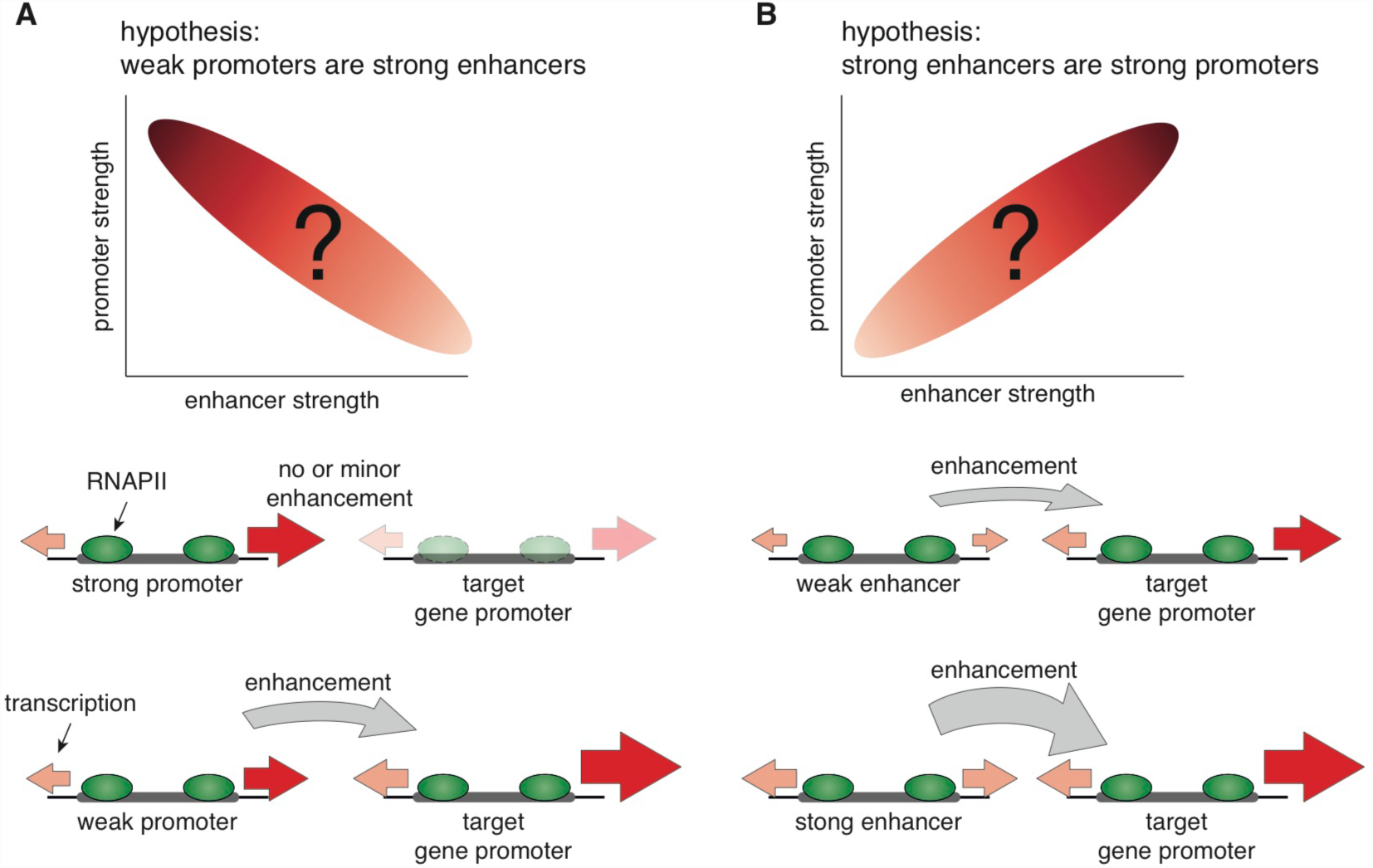
Chromatin interactions and strength of regulatory elements determine transcriptional activities. **A**: Competition between individual regulatory elements may determine their primary activity. According to this hypothesis, the individual promoter strengths of interacting gene promoters determine the enhancer activity of gene promoters according to an inverse relationship between enhancer and promoter activity. Strong gene promoters likely have no or little enhancer activities whereas weak gene promoters may function as enhancers for stronger interacting gene promoters. **B**: The transcriptional activities of enhancers and their increase in expression levels alongside with increasing expression of target gene promoters upon stimulus suggest a model in which the local transcriptional activity (promoter activity) is a proxy for its distal enhancer activity. According to this hypothesis, the strength of an enhancer is directly related to its expression level.

#### Box III: Outstanding questions

Currently, it is unclear what fraction of gene promoters also act as enhancers. Although reporter-based genome-wide assessment suggests their potential to function as enhancers [44], *in vivo* tests in which transcriptional output is measured before and after deletion of regulatory elements will be required to demonstrate this effect in genomic DNA. It is also unclear whether the ability of an element to drive or enhance transcription is static. Can weak gene promoters become stronger in certain chromatin conformations, or following the expression of a particular transcriptional activator? Can a strong promoter with little or no enhancer activity acquire enhancer activity in another context? Such flexibility may allow fine control of the spatio-temporal transcriptional activities across cells, as the function of a given regulatory element may change to fit the situation at hand. It will also be highly relevant to investigate if these mechanisms observed in mammals apply to other groups of multicellular organisms, such as insects or plants.

Many of the noted enhancer and promoter activities of regulatory elements may be influenced by confounding factors not accounted for in the study design. For instance, higher transcriptional activity of some enhancers may in some cases reflect a greater tendency to be physically proximal to RNAPII-enriched transcription factories and may therefore confound estimates of their individual promoter strengths. Thus, systematic work to decipher how these roles translate into the transcriptional activities encoded in our genomes is needed. Luckily, many of the tools needed to do this in native chromatin are becoming available, including the ability to easily edit genomic DNA with CRISPR/Cas technology [80]. Experiments leveraging these technologies to systematically delete, insert, and switch the locations of promoters and enhancers will be required to disentangle the rules governing their contextual activities *in vivo*. It will also be important to do these experiments systematically at many enhancer-promoter pairs, in order to understand the general mechanisms rather than more easily studied outliers.

## A unified architecture of regulatory elements

What determines whether RNAPII recruited by an individual regulatory element will initiate locally or distally? As outlined above, many, if not all, active enhancers have promoter activity and can initiate RNAPII. Likewise, gene promoters vary in the rates at which they recruit and initiate RNAPII, and many have the potential to enhance transcription of other gene promoters. The function of regulatory elements may further undergo contextual changes [63,64], for example serving to enhance transcription only in the proper biological conditions or chromatin context, and even the transcription of annotated gene promoters may produce unstable RNAs similar to eRNAs [40]. Thus, we believe regulatory elements should not be regarded as distinct classes with static functions but rather elements that recruit and initiate RNAPII with varying rates, and that the primary function – promoter or enhancer – is context-dependent and determined by the physical proximity of other regulatory elements and their bound factors, as well as RNA degradation signals. This view has major consequences for how transcriptional regulation is interpreted.

## Concluding remarks

The similarities between gene promoters and gene-distal enhancers in terms of their DNA sequence features, biochemical properties, and transcriptional activities strongly argue against the classical categorization of enhancers and promoters as distinct entities. Here, we suggest that gene promoters and gene-distal enhancers should be unified into a single class of regulatory element, for which the spatio-temporal activity is context-dependent and may be decided by each individual element’s promoter strength and ability to enhance transcription at other such regulatory elements. Such a unified model of regulatory elements raises a host of new questions, many of which should be addressable with current technology (Box III).

The unified architecture of regulatory elements also suggests that novel genes may efficiently arise from elements with a primary enhancer activity, as they already possess promoter potential. With rapid evolutionary changes in enhancer-flanking DNA [40], stable orphan genes may be rapidly born by mutations [65] (introduction of U1 sites or disruption of polyadenylation sites) in the sequences encoding eRNAs. Conversely, gradual insertion of polyadenylation sites downstream of a gene promoter will decrease its functional utility as a gene promoter, and only leave its potential enhancer function intact. Thus, a unified architecture of regulatory elements also provides a means to understand the evolutionary relationship between functional elements across the genome.

## Acknowledgments

This work was supported by funding from the European Research Council (ERC) under the European Union’s Horizon 2020 research and innovation programme (R.A., grant agreement No 638273), the Novo Nordisk foundation (A.S.), and the Lundbeck foundation (A.S).

